# SPOmiAlign: A modality-agnostic framework for robust and scalable spatial multimodal alignment via feature matching

**DOI:** 10.64898/2025.12.19.695434

**Authors:** Yi Wang, Zihang He, Yunjie Yan

## Abstract

Multimodal spatial omics enables systematic characterization of tissue organization by jointly profiling molecular features across transcriptomic, proteomic, and metabolomic layers. However, integrative analysis across sections and modalities is frequently hindered by non-linear tissue distortions, mismatched spatial resolutions, and the absence of shared molecular features. To address these challenges, we introduce SPOmiAlign, a modality-agnostic framework based on uncertainty-aware dense feature matching for spatial multimodal alignment. The framework achieves second-level runtime efficiency and enables accurate alignment across heterogeneous spatial omics datasets, as well as between spatial omics and histology, without manual effort or modality-specific tuning. Across diverse datasets, SPOmiAlign consistently achieves higher alignment accuracy than existing state-of-the-art methods. We further demonstrate its utility through automated registration of spatial omics data to a common coordinate framework (CCF), enabling anatomical annotation. Applying SPOmiAlign to spatial multi-omic data demonstrates how accurate multimodal alignment is essential for integrative spatial multi-omic analysis, enabling the identification of coherent spatial domains across modalities and supporting a broad range of downstream spatial analysis.

## Introduction

Multimodal spatial omics has emerged as a transformative framework for dissecting complex biological systems by concurrently capturing molecular profiles across distinct biological layers. Recent advances in spatial transcriptomics, proteomics, and metabolomics now enable direct in situ quantification of biomolecules, providing unprecedented insights into the cellular heterogeneity and intrinsic complexity of the tissue microenvironment. Crucially, spatial recording of molecular data with high-resolution histological imaging (e.g., H&E) is required to anchor molecular profiles to their precise morphological context. Although each modality contributes complementary and modality-specific information at different resolutions, the integration of multiple modalities is crucial to bridge the gap between gene expression and phenotypic execution, thus obtaining a holistic understanding of tissue architecture and function at both the cellular and molecular levels^1^.

A fundamental prerequisite for the integration of multiple spatial omics modalities is the precise spatial alignment of tissue sections. Although emerging technologies have begun enabling multimodal profiling on a single tissue section, the prevailing paradigm for multi-omics analysis still relies on data generated from consecutive or adjacent sections. This physical separation introduces substantial non-linear geometric distortions and morphological inconsistencies due to tissue processing, coupled with discrepancies in spatial resolution across modalities. Therefore, accurate alignment of multimodal spatial omics sections is essential to faithfully integrate heterogeneous molecular and morphological information, allowing meaningful comparisons that reveal true spatial relationships^2^. Furthermore, the registration of spatial omics data in a Common Coordinate Framework (CCF) extends the utility by facilitating standardized anatomical referencing across samples and studies, which is critical for cross-dataset integration and large-scale comparative analyses^3^.

A major bottleneck in spatial multimodal alignment is the substantial heterogeneity of the data in distinct modalities. Unlike unimodal alignment, where sections share similar contrast mechanisms, spatial omics datasets encompass disparate data distributions—ranging from sparse, discrete gene counts (transcriptomics) and continuous spectral intensities (metabolomics/proteomics) to dense, high-resolution optical densities (histology). These modalities exhibit fundamentally different signal-to-noise profiles, dynamic ranges, and spatial resolutions, resulting in a lack of direct visual or statistical correspondence. Consequently, standard alignment algorithms often fail to reconcile these incommensurable feature representations, making it difficult to perform cross-modal spatial alignment.

Current spatial alignment methodologies can be generally categorized into image-to-image^4–7^, spatial multi-omics^8–20^, and spatial omics-to-image alignment^21,22^. However, a unified framework capable of seamlessly generalizing across these diverse scenarios remains unavailable, particularly one that can robustly align arbitrary spatial modalities or anchor molecular data to histological references without modality-specific tuning. Moreover, computational scalability poses a significant bottleneck; many existing tools suffer from substantial computational overhead, limiting their scalability to high-resolution, atlas-scale datasets. In addition, current approaches are predominantly based on manually annotated landmarks^3^, making the process labor-intensive, time-consuming, compromising reproducibility, and limiting the feasibility of large-scale studies.

Here, we propose a modality and technology-agnostic framework for spatial multimodal section alignment, capable of registering heterogeneous spatial modalities and datasets generated by different technologies within the same modality. Extensive benchmarking demonstrates that SPOmiAlign achieves superior alignment accuracy while significantly reducing computational overhead. Notably, our framework enables the fully automated registration of spatial omics data onto a CCF, allowing the rapid retrieval of standardized CCF anatomical annotations. By applying SPOmiAlign to mouse brain multimodal datasets, including spatial transcriptomics, spatial proteomics, and spatial metabolomics, we demonstrate that the accurate alignment achieved by SPOmiAlign facilitates integrative spatial multi-omics analysis, enabling the identification of previously unrecognized functional spatial domains defined by coherent multi-omic signatures.

## Results

### Overview of SPOmiAlign

To enable robust spatial alignment across heterogeneous spatial omics modalities and experimental technologies, we applied SPOmiAlign to multimodal spatial datasets derived from adjacent tissue sections as well as from independent specimens. This modality- and technology-agnostic framework allowed consistent spatial registration across datasets, supporting downstream analyses in both spatial omic-to-image and spatial omic-to-omic integration contexts (Fig. 1**a**).

**Figure 1.**
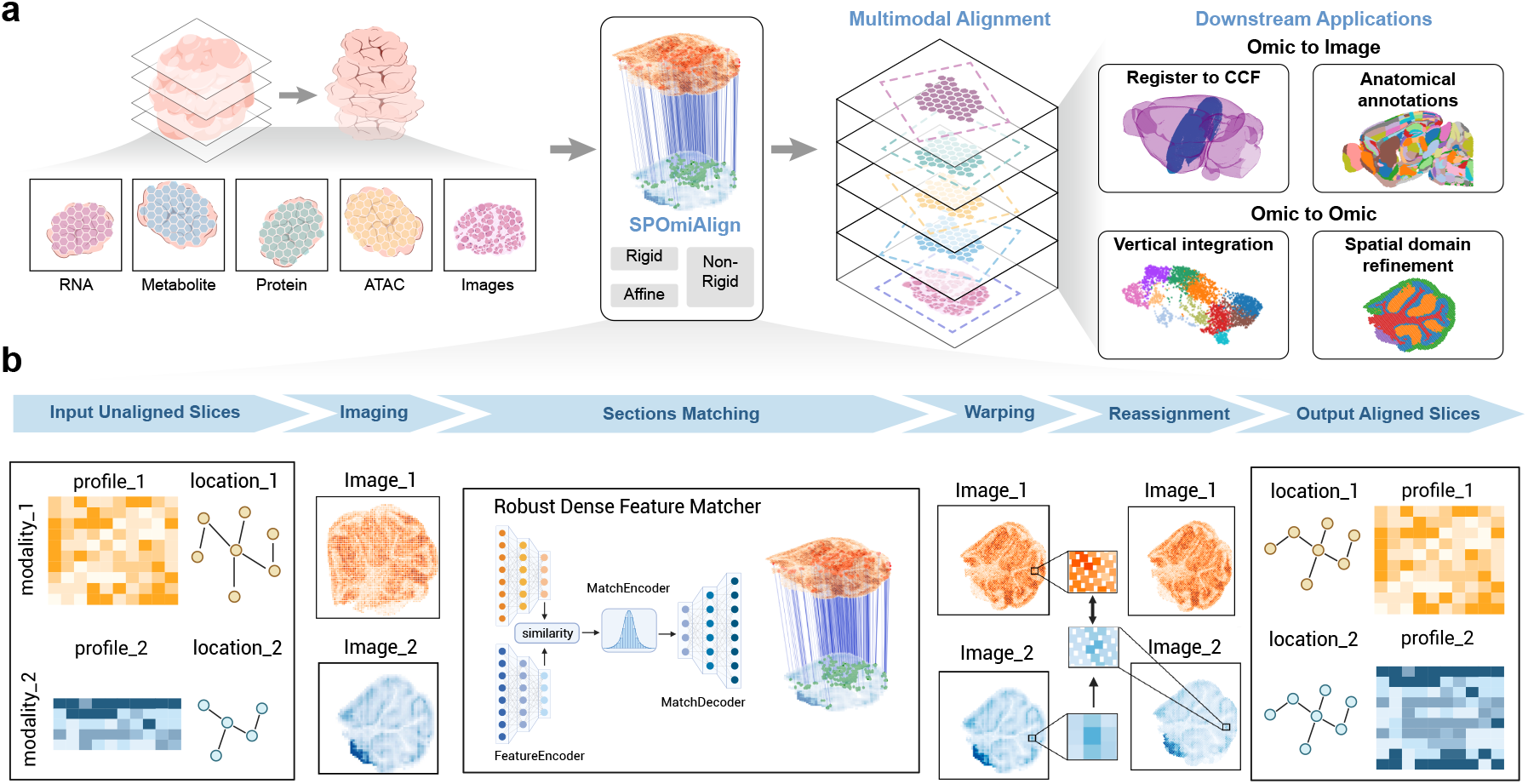
Overview of SPOmiAlign for modality-agnostic spatial multimodal alignment. **a** End-to-end application workflow. Spatial multi-omics profiles are generated from adjacent tissue sections using complementary technologies. SPOmiAlign performs spot-level alignment across modalities and sections, producing aligned datasets that support downstream tasks in two major settings: omic-to-image alignment (e.g., mapping to histology or an anatomical atlas/CCF) and omic-to-omic alignment for vertical multimodal integration. **b** Algorithmic pipeline of SPOmiAlign. For omics inputs, each section (reference and source) is provided as spot-level molecular profiles with spatial coordinates and is rendered into a spatial structural image (SSI) for image-based matching; for image-modal inputs, the reference and source images are used directly. The section matching module comprises a DINOv2-based feature encoder, a Gaussian process (GP)-based matching encoder, and a Transformer-based matching decoder, producing dense pixel-wise correspondences with confidence scores. In the section warping module, a high-confidence correspondence subset (top 5,000 matches) is used to estimate a geometric transformation, which is then applied to warp the source section (and its spot coordinates). For omic-to-omic alignment, a reassignment step establishes one-to-one spot correspondence between modalities after warping, enabling downstream integration and comparative analysis.

In the spatial omic-to-image aspect, SPOmiAlign enabled the reliable registration of spatial omics sections onto a common coordinate framework (CCF), such as the Allen Brain Atlas^23^, which allows the automated retrieval of annotations from the anatomical region and facilitates region-based downstream analyses. In the spatial omic-to-omic aspect, it facilitates vertical integration and spatial domain refinement by ensuring spatial correspondence across modalities.

The SPOmiAlign framework consists of four major components: imaging, section matching, warping, and reassignment (Fig. 1**b**). In the imaging module, spatial omics profiles are converted into image-like representations to enable cross-modality comparison based on spatial structural features. For spatial omics sections with paired imaging data, including histological or fluorescence images, the paired image data were used for downstream alignment. For spatial omics sections lacking paired images, molecular profiles were transformed into spatial structural images (SSIs) that capture spatial organization and enable subsequent image-based registration. Here, grayscale SSIs were constructed by aggregating gene or feature profiles per spatial spot, so that the intensity of the pixels reflects the overall expression or abundance at each location. Alternatively, high-dimensional omics profiles can be projected into low-dimensional spaces via principal components or cell-type annotations and encoded as RGB images to preserve richer spot information.

To align two image-based sections, SPOmiAlign first identified dense correspondences between a reference section and a source section using a robust section-matching strategy. Unlike approaches relying on manually annotated landmarks, SPOmiAlign leverages dense feature matching via RoMa^24^, a pretrained correspondence model optimized for diverse imaging domains. By inferring dense point-wise correspondences across entire tissue sections, this strategy provides stable performance across heterogeneous modalities and spatial resolutions while substantially reducing computational overhead compared to iterative optimization-based or landmark-driven alignment schemes^15^. Based on the matched correspondences, geometric transformations were estimated and applied through a dedicated warping module, supporting rigid, affine, and non-rigid transformations to correct both global orientation differences and local geometric distortions, thereby enabling scalable alignment of large spatial omics datasets.

Because differences in spatial resolution and sampling densities across modalities often prevent a one-to-one correspondence between spots even after geometric alignment, SPOmiAlign incorporates a reassignment module to reconcile these discrepancies. Using a nearest-neighbor strategy, each spot in the warped source section is reassigned to its closest spatial neighbor in the reference section, enabling one-to-one spot correspondence across sections. This reassignment step is particularly important for vertical downstream integration that requires spatially matched observations across datasets.

### SPOmiAlign enables automated spatial omic-to-CCF registration and anatomical annotation retrieval

Accurate registration of spatial omics data in a common coordinate framework (CCF) is critical for interpreting molecular patterns within their anatomical context. However, most existing approaches largely rely on manual or semi-manual landmark selection, limiting scalability across datasets and technologies^3,25^. Using its ability to perform omic-to-image alignment, SPOmiAlign enables fully automated registration of two-dimensional sections in an anatomical atlas, such as the Allen Brain Atlas, thereby supporting standardized anatomical annotation at single-spot resolution.

Specifically, SPOmiAlign maps experimental spatial transcriptomics sections to the Allen Brain Atlas by first identifying the Nissl-stained image with the most anatomically corresponding from the atlas and aligning the experimental section to this reference (Fig. 2**a**). Following registration, each spatial spot is assigned to a CCF coordinate, from which standardized anatomical annotations are retrieved.

**Figure 2.**
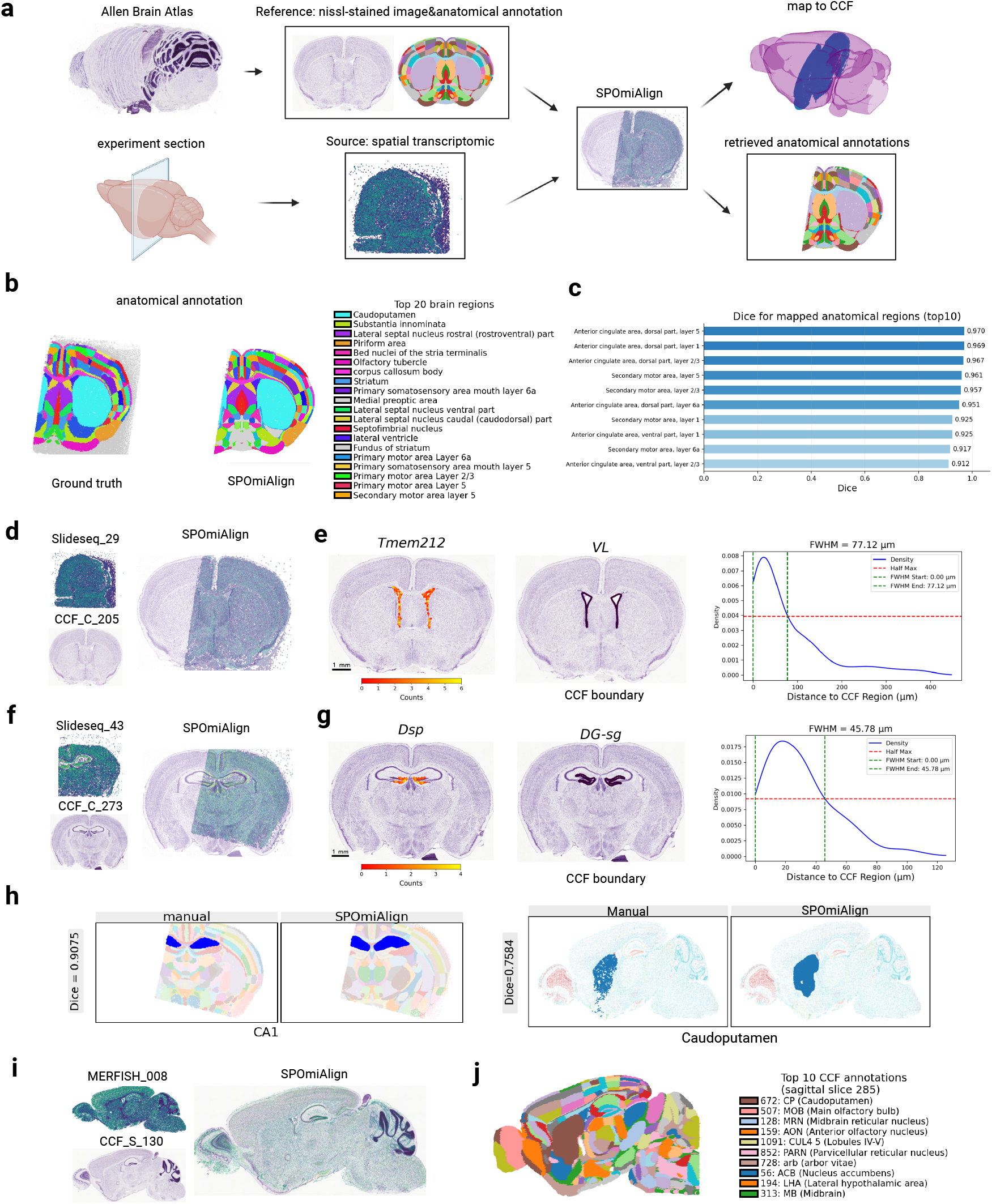
SPOmiAlign enables automated spatial omic-to-CCF registration and anatomical annotation retrieval. **a** Schematic overview of automated spatial omics registration to a CCF using SPOmiAlign. **b** Comparison of anatomical annotations obtained by SPOmiAlign and ground truth, with the top ten brain regions ranked by area shown in the legend. **c** Dice similarity coefficients for the top ten brain regions with the highest annotation agreement between SPOmiAlign and ground truth. **d, f** Overlay of Slide-seq mouse brain coronal sections (IDs 29 and 43) with matched CCF Nissl-stained images after alignment by SPOmiAlign. **e, g** Distance density distributions between enriched gene expression spots and corresponding CCF region boundaries for Slide-seq sections 29 and 43. Full width at half maximum (FWHM) quantifies the spatial deviation of molecular signals from anatomical boundaries. **h** Dice coefficient comparison between SPOmiAlign and ground truth annotations for representative regions: CA1 in Slide-seq section 43 (left) and Caudoputamen in MERFISH sagittal section 008 (right). **i** Overlay of MERFISH mouse brain sagittal section 008 with the matched CCF reference section after alignment. **J** Anatomical annotation of MERFISH section 008 assigned by SPOmiAlign, with the top ten regions by area shown in the legend (CCF annotation section at *z* = 285).

To systematically evaluate alignment performance across different spatial transcriptomics technologies, we analyzed two three-dimensional mouse brain datasets generated using distinct platforms: a sequencing-based Slide-seq coronal dataset and an imaging-based MERFISH sagittal dataset^3,25^. Representative two-dimensional sections exhibiting clear anatomical structures were selected for evaluation. For Slide-seq section IDs 29, visual inspection confirmed strong spatial concordance between the retrieved Nissl-stained images and the aligned spatial transcriptomics sections (Fig. 2**d,f**, ID 43 in Supplementary Fig. 1b).

For the Slide-seq dataset, anatomical annotations from Allen Brain Atlas(Method) were used as ground truth to benchmark the anatomical labels generated by SPOmiAlign. At a qualitative level, the automatically assigned region annotations closely recapitulated the spatial organization of the manually curated labels across ID 29 coronal sections (Fig. 2**b**, Supplementary Fig. 1b). The registration accuracy was quantified using the Dice coefficient, which measures the spatial overlap between the corresponding anatomical region masks. Across the top ten regions with the highest alignment accuracy, the Dice coefficients exceeded 0.91 for Slide-seq section ID 29 (Fig. 2**c**, Supplementary Fig. 1c), with particularly strong concordance observed in fine-grained cortical regions such as the anterior cingulate area and the secondary motor area, highlighting the precision of SPOmiAlign in aligning anatomically detailed structures.

To assess biological concordance beyond geometric overlap, we examined genes with well-established region-specific expression patterns. Consistent with manual annotations, *Dsp* was enriched in the dentate gyrus subgranular zone (DG-sg), while *Tmem212* showed preferential expression in the ventral lateral (VL) thalamic region. For each gene, we quantified the spatial deviation between high-expression spots and atlas-defined region boundaries by measuring the full width at half maximum (FWHM) of the distance distribution^3^. Across enriched genes, the FWHM values ranged from 40 to 70 *µ*m, comparable to the precision typically achieved by expert manual alignment (Fig. 2**e, g**, Supplementary Fig. 1d), indicating that region-specific molecular signals remain spatially coherent after SPOmiAlign mapping. Representative regions, including the cornu ammonis area 1 (CA1, Slide-seq ID 43, Fig. 2**h**, caudoputamen and piriform cortex (Slide-seq ID 29; Supplementary Fig. 1a), each achieved Dice coefficients greater than 0.72.

In addition, we evaluated SPOmiAlign on the MERFISH sagittal dataset. Despite substantial differences in measurement modality, imaging resolution, and spatial organization, SPOmiAlign produced coherent anatomical mappings (Fig. 2**i, j**). The dice scores for representative regions (Fig. 2**h**) demonstrated high agreement with the atlas annotations, underscoring the robustness of SPOmiAlign across different technologies.

### SPOmiAlign enables robust spatial omic-to-omic alignment across samples via histology-guided registration

To evaluate the performance of SPOmiAlign for spatial omic-to-omic alignment across samples, we analyzed a multi-omic mouse brain dataset consisting of spatial transcriptomics (sample 1, S1) and spatial ATAC-seq (sample 2, S2) generated from two independent mice^26^. Because spatial alignment between samples is frequently confounded by tissue deformation introduced during sectioning and processing, we further applied synthetic geometric perturbations, including rotation, translation, and scaling, to the spatial coordinates of S2 to systematically assess alignment robustness. As each section was associated with a paired H&E histology image, these images were used as intermediate references to allow image-guided alignment between the two spatial omics sections.

Paired spatial structural images (SSIs) and H&E-stained sections from samples S1 and S2 are shown prior to alignment (Fig. 3a). SPOmiAlign was compared with ELD^15^, an automated multimodal image registration method, and STAlign^14^, a spatial transcriptomics alignment approach, as well as with unaligned sections. Global overlap analysis showed that SPOmiAlign achieved qualitatively accurate alignment, particularly in regions with weak structural features such as tissue boundaries (Fig. 3b).

**Figure 3.**
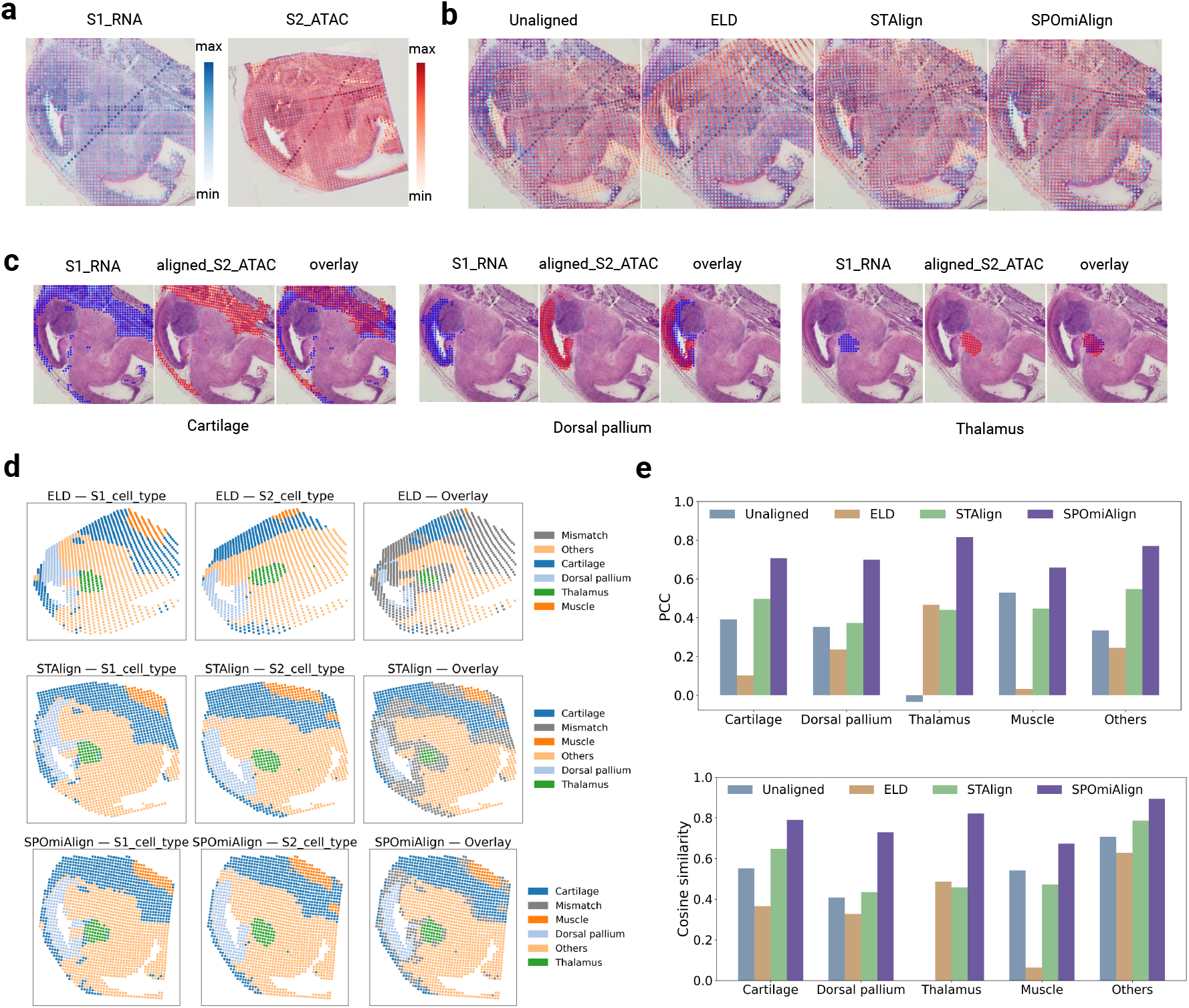
SPOmiAlign enables robust spatial omic-to-omic alignment across samples via histology-guided registration. **a** MISAR-seq mouse brain spatial bi-omics dataset, where Sample 1 (S1) and Sample 2 (S2) were profiled by spatial transcriptomics and spatial ATAC-seq, respectively. Spatial structural images (SSIs) are overlaid with paired H&E-stained histology images. **b** SSI overlays of S1 and S2 after alignment using SPOmiAlign, ELD, STAlign, and without alignment. **c** Overlap of representative cell types between aligned S2 and S1 following SPOmiAlign. **d** Visualization of cell-type annotations for S1 and aligned S2 under different alignment methods; black squares indicate mismatched spots between corresponding cell types. **e** Quantitative evaluation of spatial concordance between S1 and S2 across cell types using Pearson correlation coefficient (PCC) and cosine similarity for different alignment methods.

For quantitative evaluation, we adopted spatial correlation metrics at the cell-type level, including the Pearson correlation coefficient (PCC) and cosine similarity. Manual cell-type annotations from the spatial transcriptomics section (S1) and clustering-based annotations from the ATAC-seq section (S2) were harmonized to ensure correspondence across datasets, with unmatched labels grouped into a category” other. Visual inspection of representative cell types revealed qualitatively accurate overlap after alignment with SPOmiAlign (Fig. 3c). Consistently, the number of mismatch spots, defined as spatially inconsistent cell-type assignments, was markedly reduced compared to both ELD and STAlign (Fig. 3d). Across the corresponding cell types, SPOmiAlign achieved the highest spatial concordance, with values of PCC and cosine similarity predominantly ranging from 0.7 to 0.8.

To assess scalability for high-resolution spatial datasets, we further evaluated computational efficiency by repeating the alignment procedure five times using SPOmiAlign and STAlign. Runtime analysis demonstrated that SPOmiAlign consistently achieves alignment in 10 seconds. In contrast to STAlign, which typically requires minutes to hours for alignment, SPOmiAlign reduces computational time by two to three orders of magnitude (Supplementary Fig. 2). This substantial speedup highlights the scalability of SPOmiAlign and underscores its suitability for future spatial omics datasets with rapidly increasing spatial resolution.

To further validate the non-rigid image-to-image registration component underlying SPOmiAlign, we evaluated its performance on the ANHIR benchmark dataset, a public reference standard for non-rigid histological image registration^27^. ANHIR comprises serial or adjacent histological sections exhibiting substantial non-linear deformations and staining variability. Across diverse tissue types, disease conditions, and staining protocols, SPOmiAlign consistently achieved accurate and efficient alignment, demonstrating robust performance under challenging non-rigid distortions (Supplementary Fig. 3). Together, these results demonstrate that SPOmiAlign achieves competitive performance for non-rigid image-to-image registration, in addition to its effectiveness in image-guided spatial omic-to-omic alignment.

### SPOmiAlign supports trimodal spatial omic-to-omic alignment and facilitates spatial multi-omic integration analysis

Recent advances in spatial multi-omics have expanded beyond bimodal analyses toward trimodal and higher-order profiling, enabling more comprehensive characterization of tissue organization by jointly capturing transcriptional, proteomic, and metabolic states. However, increasing the complexity of the modality introduces substantially greater challenges for accurate spatial alignment across datasets, particularly in the absence of paired histological or imaging references.

To assess trimodal alignment performance, SPOmiAlign was evaluated on a mouse brain spatial trimodal dataset comprising three adjacent tissue sections profiled using distinct technologies: spatial transcriptomics generated by MAGIC-seq, spatial proteomics measured by PLATO, and spatial metabolomics acquired via MALDI–MSI^28^. The spatial transcriptomics section served as the reference, while the spatial proteomics and metabolomics sections independently aligned with the transcriptomic coordinate system. In the absence of paired imaging data, all spatial omics sections were converted to spatial structural images (SSIs) for alignment. Before alignment, pronounced spatial discrepancies were observed between the SSIs of the three modalities, most notably in the spatial metabolomics section, which exhibited substantial positional offsets relative to the other two modalities (Fig. 4a). Consequently, an initial manual transformation (rotation, translation, and scaling; referred to as manual) was applied to the metabolomics section before alignment, serving as a common preprocessing step for all the compared methods.

**Figure 4.**
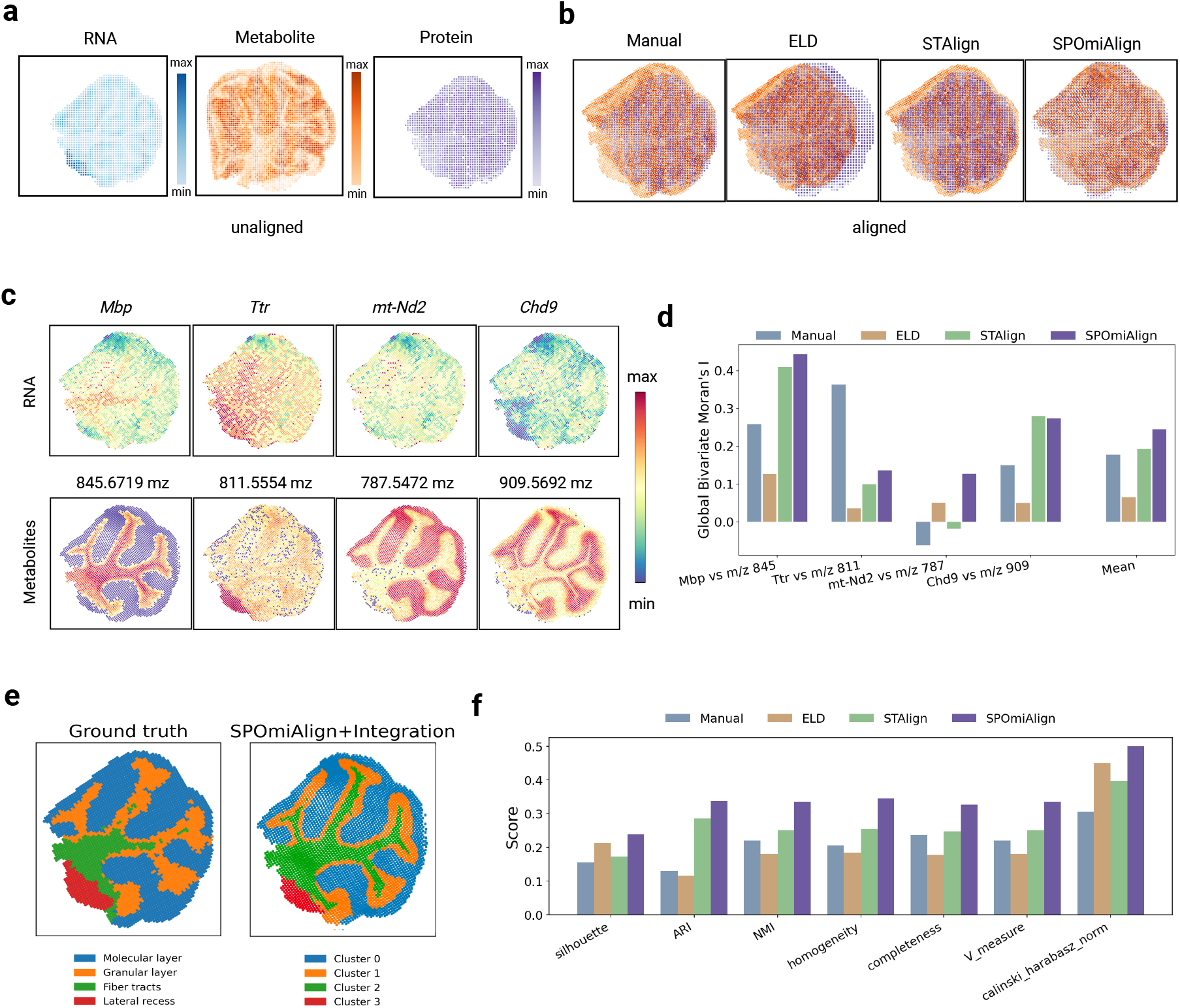
SPOmiAlign supports trimodal spatial omic-to-omic alignment and facilitates spatial multi-omic integration. **a** Spatial structural images (SSIs) of three adjacent mouse brain sections profiled by spatial transcriptomics, spatial proteomics, and spatial metabolomics before alignment. **b** Overlay of aligned SSIs produced by SPOmiAlign, STAlign, ELD, and manual affine pre-alignment, illustrating trimodal spatial correspondence after alignment. **c** Spatial distribution heatmaps of four pairs of spatially co-localized biomarkers across spatial transcriptomics and spatial metabolomics modalities. **d** Bivariate Moran’s *I* values for the four biomarker pairs after alignment by SPOmiAlign, STAlign, ELD, and manual alignment, together with the mean Moran’s *I* for each method.

Benchmarking against STAlign, ELD, and the manually transformed baseline demonstrated that SPOmiAlign achieved consistently improved global alignment, with enhanced correspondence at tissue boundaries and finer local anatomical structures (Fig. 4b). Quantitative assessment was performed using four pairs of spatially co-localized biomarkers previously reported to exhibit consistent spatial distributions across spatial transcriptomics and spatial metabolomics modalities (Fig. 4c). The precision of the alignment was quantified using the bilinear Moran’s I statistic, which measures spatial co-localization between two variables. Across all biomarker pairs, SPOmiAlign yielded the highest mean bilinear Moran’s I values (Fig. 4d), indicating superior spatial concordance relative to alternative methods.

To assess whether improved alignment results in improved downstream multimodal integration, integrative analysis was performed on the aligned trimodal dataset using SpatialGLUE^29^. Because spatial transcriptomics data included cell-type annotations, integration quality was evaluated by comparing clustering results with the original transcriptomic annotations. When clustering into four domains, the integrated data exhibited strong concordance with known cell-type annotations, while additionally revealing fiber tract structures that were not clearly identifiable in the unimodal transcriptomics (Fig. 4e). Across multiple standard integration quality metrics, including normalized mutual information (NMI) and adjusted Rand index (ARI), SPOmiAlign-based alignment consistently outperformed STAlign, ELD and manually aligned datasets, demonstrating that accurate spatial alignment is critical for high-quality trimodal multimodal integration (Fig. 4f). Together, these results demonstrate that precise spatial alignment is a critical prerequisite for effective spatial multi-omic integration, highlighting the importance of robust spatial alignment frameworks such as SPOmiAlign for advancing spatial multi-omic analysis.

### SPOmiAlign enables the discovery of fine-grained molecular stratification within the cerebellar molecular layer

Building on the alignment and integration benchmarks above, we next evaluated whether SPOmiAlign could resolve substructures within the cerebellar molecular layer that were not apparent under the original annotation. The partition comprising five groups was supported by consistent cluster connectivity across adjacent resolutions in Clustree and by concordant modality-specific signals after alignment (Fig. 5a, Supplementary Fig. 4a)^30^. At this resolution, the molecular layer was subdivided reproducibly into two spatial sublayers, here termed the inner molecular layer (IML) and the outer molecular layer (OML), while other anatomical regions, including the granular layer, fiber tracts, and lateral recess, remained consistent with the original annotation^28^. This subdivision was robust across adjacent resolutions and supported by concordant patterns across multiple molecular modalities.

**Figure 5.**
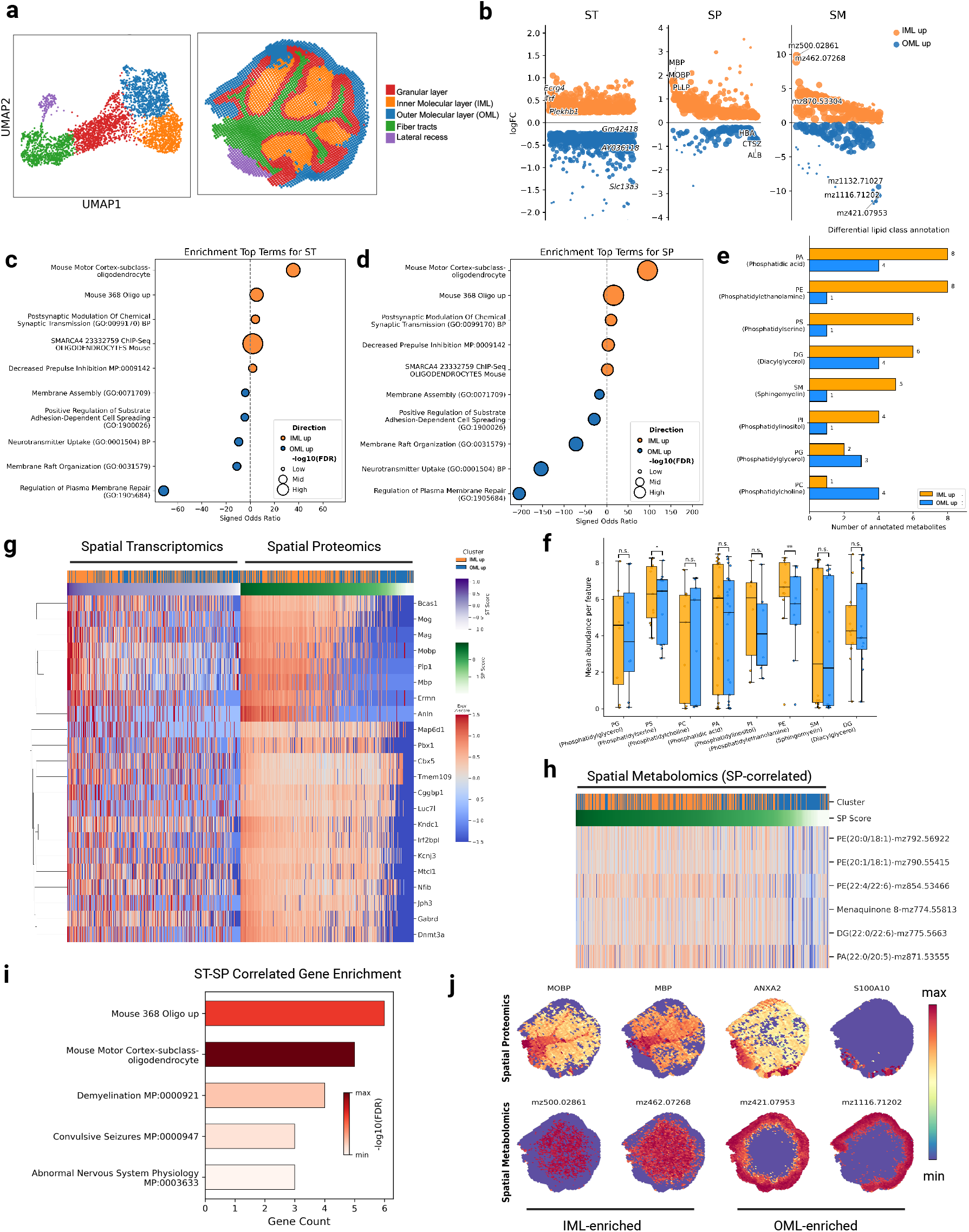
Spatial multi-omics dissection of molecular layer substructure in the mouse cerebellum. **a** Identification of stable Inner (IML) and Outer (OML) Molecular Layer subdivisions. **b** Differential analysis of ST, SP, and SM features between sublayers (logFC *>* 0.25, FDR < 0.1). **c**,**d** Shared pathway enrichment for ST (**c**) and SP (**d**), highlighting divergent biological programs (FDR *<* 0.05). **e** Differential lipid class annotation based on spatially variable metabolites (SVM), showing the distribution of annotated lipid classes. **f** Mean intensity quantification of lipid classes (paired Student’s *t* test; * *P*< 0.05, ** *P*< 0.01, n.s., not significant). **g** Heatmap of jointly regulated spatially variable gene–protein pairs (SVG–SVP) identified from integrated ST–SP analysis. Features are ordered by decreasing mean expression across spatial spots. Only positively correlated pairs are shown (Spearman’s *ρ >* 0.1, FDR *<* 0.05). **h** Heatmap of annotated metabolites correlated with protein signatures from SVG–SVP concordant pairs (Spearman’s *ρ >* 0.2, FDR *<* 0.05). **i** Functional enrichment analysis of concordant SVG–SVP signatures (FDR *<* 0.05). **j** Spatial visualization of representative proteins and metabolites identified from integrated multi-omics analyses confirming sublayer specificity. Left: IML-enriched features; Right: OML-enriched features.

Differential analysis of spatial transcriptomics (ST) and spatial proteomics (SP), together with enrichment analysis restricted to pathways detected in both modalities, revealed marked molecular divergence between the two sublayers (Fig. 5b–d). The IML showed a strong enrichment of myelin-associated genes and proteins, including *MOBP* and *MBP*, as well as oligodendrocyte-associated programs, consistent with the established roles of mature oligodendrocytes in the maintenance and axonal support (Fig. 5j, Supplementary Fig. 5a) ^31,32^. Concurrently, the IML exhibited elevated expression of neurofilament proteins (*NEFH, NEFL, NEFM*; Supplementary Fig. 5b). Given that neurofilaments are ubiquitous structural components of neuronal cytoskeletons, their higher signal in the IML is consistent with an increased local axonal content. Together with oligodendrocyte-enriched transcriptional and proteomic programs, this pattern supports a spatial association between axon-rich compartments and glial programs within the deep molecular layer^33^.

In contrast, OML was characterized by the enrichment of pathways related to membrane organization, assembly, and lipid turnover, including annexin-associated processes involving *ANXA2* and *S100A10* (Fig. 5c,d,j). These pathways have been implicated in calcium-dependent membrane dynamics, cytoskeletal remodeling, and vesicular trafficking^34,35^. The OML also showed elevated expression of plasma-associated proteins (*ALB, HBA*) and the Purkinje-related marker *PCP4L1*, consistent with its anatomical proximity to Purkinje cell dendrites and local vasculature (Supplementary Fig. 5c) ^36^. Collectively, these observations point to a compartment defined by increased baseline membrane turnover and synaptic activity. Although these signatures are compatible with a membrane-dynamic compartment near synaptic and perivascular structures, they may also reflect high baseline trafficking and lipid metabolism in active dendritic microenvironments.

Spatial metabolomics (SM) analysis revealed distinct lipid stratification consistent with the subregional organization of the molecular layer. Although the absolute number of differentially annotated lipid classes varied between sublayers (Fig. 5e), quantitative analysis of the mean abundance of features revealed a specific enrichment of myelin-associated lipids within IML (Fig. 5f). Specifically, phosphatidylethanolamine (PE) and phosphatidylserine (PS) were markedly elevated in IML compared to OML. PE is abundant in myelin membranes and contributes to membrane curvature and packing, while PS, although typically a minor fraction, may influence membrane asymmetry and protein–lipid interactions^34,37,38^. It is important to note that the annotation of metabolites based solely on *MALDI-MSI m/z* matching is inherently limited in its ability to resolve isomers or strictly validate chemical identities. Therefore, these findings should be interpreted as reflecting the collective spatial trends of lipid classes, subject to the intrinsic constraints of the annotation resolution of the dataset^28^.

In the integrated multi-omics landscape, the synergy between modalities further refined this stratification. Although joint ST–SP enrichment analysis successfully identified shared myelin and oligodendrocyte-related pathways in IML, the spatial correlation heatmap revealed distinct modality-specific characteristics: spatial proteomics (SP) exhibited significantly clearer cluster-specific patterning (IML versus OML) compared to spatial transcriptomics (ST), which showed more diffuse spatial gradients (Fig. 5g,i). Despite modest spot-level correlations (Supplementary Figure 5g,h), multiple lipid species showed reproducible, direction-consistent associations with IML-enriched RNA or proteins between spots (Supplementary Figure 5d–f). The SP–SM correlation analysis identified specific lipid species, including multiple PE molecules, which were strongly tracked with the proteomic signature of IML (Fig. 5h).

Together, these results validate the biological coherence of the identified sublayers and provide an integrated view of tissue organization that covers transcriptional programs, protein expression, and metabolic state.

In addition, based on the clustree analysis, we identified two additional stable clustering resolutions at higher granularity, corresponding to cluster numbers of 7 and 12. For each resolution, we visualized the resulting spatial domains together with their corresponding UMAP embeddings. These visualizations suggest that SPOmiAlign-based alignment has the potential to improve the refinement and delineation of spatial domains (Supplementary Fig. 5b).

## Discussion

In this study, we present SPOmiAlign, a modality- and technology-agnostic spatial alignment framework that enables accurate and scalable alignment across heterogeneous spatial multi-omic datasets and between spatial omics and histological images.

Building on robust dense feature matching, uncertainty-aware correspondence modeling, and coarse-to-fine geometric refinement, SPOmiAlign overcomes key challenges associated with non-rigid tissue deformation, mismatched spatial resolution, partial tissue overlap between sections, and the absence of shared molecular features. Across a broad range of alignment scenarios, including spatial omics-to-CCF registration, spatial omics-to-omics alignment, and trimodal spatial integration, SPOmiAlign consistently achieved high alignment accuracy while substantially reducing computational cost. Together, downstream biological analyses demonstrate that spatial alignment is a foundational determinant of biological interpretability in spatial multi-omics studies.

A central strength of SPOmiAlign lies in its modality-agnostic design, which decouples spatial alignment from modality-specific molecular features. By transforming spatial omics data into image-based representations or directly leveraging paired imaging data when available, SPOmiAlign enables a unified alignment strategy across sequencing-based, imaging-based, and mass spectrometry-based spatial technologies. This contrasts with many existing methods that rely on shared gene expression features, manual landmark annotation, or modality-specific assumptions, which often limit generalizability and scalability. By abstracting alignment to a common structural representation, SPOmiAlign allows spatial correspondence to be established independently of the molecular layer under investigation, substantially broadening its applicability.

Another key advantage of SPOmiAlign is the use of robust dense feature matching for section correspondence. Using a pretrained dense correspondence foundation model, SPOmiAlign infers dense, uncertainty-aware correspondences across entire tissue sections in a single forward pass, rather than relying on iterative optimization over sparse landmarks. This strategy improves robustness to strong non-rigid distortions and weakly textured regions, such as tissue boundaries, providing a reliable basis for precise anatomical annotation and high-quality spatial multi-omics integration. Importantly, dense correspondence inference enables anatomically faithful alignment even in regions where molecular signals are sparse or modality-specific, a scenario commonly encountered in real-world spatial omics datasets.

SPOmiAlign is further distinguished by its computational efficiency. Dense correspondence inference followed by lightweight geometric estimation allows alignment to be completed in seconds, avoiding the iterative optimization procedures that characterize many existing alignment tools. Compared with alternative methods, which typically require minutes to hours, SPOmiAlign reduces runtime by two to three orders of magnitude. This efficiency enables scalable applications to high-resolution, atlas-scale datasets and facilitates exploratory or iterative analyses that would otherwise be computationally prohibitive.

Beyond methodological performance, SPOmiAlign provides substantial biological and analytical utility by enabling anatomical standardization of spatial omics data. Automated registration of spatial omics data to a common coordinate framework enables standardized anatomical annotation, allowing molecular patterns to be interpreted within a shared anatomical reference across samples, technologies, and studies. This capability is particularly valuable for large-scale atlas projects and cross-dataset comparisons, where manual annotation is impractical and prone to inconsistency. By embedding spatial omics data within a unified anatomical context, SPOmiAlign supports reproducible, region-aware analyses that extend beyond individual experiments.

Moreover, accurate spatial alignment is a critical prerequisite for integrative spatial multi-omics analysis. Our results demonstrate that improved alignment directly enhances multimodal integration, enabling the identification of coherent spatial domains defined by consistent transcriptional, proteomic, and metabolic signatures. In particular, SPOmiAlign-facilitated integration revealed fine-grained spatial substructures that are typically obscured by misalignment, underscoring the importance of precise spatial correspondence for uncovering biologically meaningful patterns.

Despite the strong performance of SPOmiAlign, several limitations remain. When paired imaging data are unavailable, spatial structural images (SSIs) are constructed from discrete spots with manually specified sizes to approximate pixel continuity. Because spatial resolution varies across modalities, this manual choice can introduce user intervention and influence alignment outcomes. Future work could mitigate this limitation by incorporating interpolation-based approaches, such as adaptive kernel density estimation or spatial interpolation. In addition, the current SSI formulation aggregates high-dimensional molecular features into a single scalar intensity, which prioritizes global structural contrast but may attenuate fine-grained molecular heterogeneity or modality-specific signals. This abstraction reflects an inherent trade-off between geometric robustness and molecular specificity and does not aim to preserve the full informational content of the original omics profiles. In future studies, we will focus on designing tissue- or modality-adaptive SSI constructions, including multi-channel or feature-weighted representations, to better capture context-specific structural cues.

In summary, SPOmiAlign provides a robust, scalable, and computationally efficient framework for spatial alignment across diverse spatial omics modalities and technologies. As spatial multi-omics datasets continue to grow in resolution, complexity, and scale, SPOmiAlign offers a practical solution for unifying spatial information across molecular layers and experimental platforms and facilitates a broad range of integrative spatial multi-omic studies.

## Methods

### Data Preprocessing in SPOmiAlign

#### Construction of Spatial Structural Images (SSIs)

To enable a unified and spatially comparable representation of spatial multi-omics datasets, we transform multimodal molecular profiles into a common image-based form that preserves underlying tissue architecture. Specifically, we construct a Spatial Structural Image (SSI) by aggregating multi-dimensional molecular features at each spatial spot into a single intensity value, thereby converting high-dimensional spatial omics measurements into a single-channel image representation. Formally, a spatial omics sample consists of *N* spatial locations (spots), indexed by *i* = 1, …, *N*. At each location *i*, we define a feature vector of *D*-dimensional.

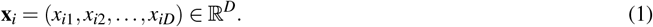

SSI is a scalar-valued image obtained by applying a feature aggregation function *f* (·) to each spot:

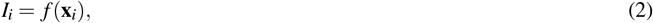

where *I*_*i*_ denotes the intensity of SSI at the point *i*. We adopt summation over features:

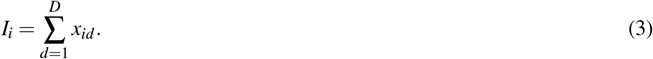

Finally, SSI is rendered by assigning *I*_*i*_ to the spatial coordinate (*u*_*i*_, *v*_*i*_) of the spot *i*, producing a two-dimensional single-channel image that preserves the spatial organization of the tissue.

#### Registering Slide-seq mouse brain coronal sections to the Allen Brain Atlas

We used the Slide-seq mouse brain spatial transcriptomics dataset comprising 101 coronal sections spanning the anteroposterior axis of the adult mouse brain. Following the original study, we selected two representative sections (sections 29 and 43) for registration in the Allen Brain Atlas. For each selected section, we constructed the corresponding SSI as described above. To enhance anatomical contrast in SSIs, we filtered spatial spots by RNA abundance and retained only spots whose UMI counts were within the top 20% of the section (i.e., above the 80th percentile of the UMI distribution). The original Slide-seq data have a pixel resolution of 1.3 *µ*m. During the SSI rendering, each retained spot was drawn as a filled disk with a radius of 5 pixels, corresponding to an effective spatial coverage of approximately 13 *µ*m per spot.

#### Registering MERFISH sagittal mouse brain sections to the Allen Brain Atlas

To evaluate the applicability of SPOmiAlign on mouse brain sagittal sections and imaging-based spatial transcriptomics, we registered a representative MERFISH sagittal section (midline section 008 among 25 sagittal sections) to the Allen Brain Atlas. We first constructed the SSI for the selected MERFISH section. During SSI rendering, each spatial spot was visualized at twice the original MERFISH spatial resolution.

#### Anatomical annotation from the Allen Brain Atlas

The Allen Brain Atlas provides high-resolution anatomical reference images together with a systematic and hierarchical taxonomy of mouse brain structures. In this study, we used the coronal atlas (132 coronal sections spaced at 100 *µ*m intervals) and the sagittal atlas (21 sagittal sections spaced at 200 *µ*m intervals). Each atlas includes Nissl-stained images and corresponding anatomical annotations. For annotation retrieval, the required inputs include the index of the CCF reference section corresponding to the experimental slice and an anatomical annotation volume. Specifically, Slide-seq coronal sections 29 and 43 were mapped to CCF coronal sections 205 and 273, respectively, while MERFISH sagittal section 008 was mapped to CCF sagittal section 130 by manual inspection. Anatomical annotations were obtained from the Allen CCF 2017 release, which provides annotation volumes at multiple isotropic resolutions (10, 25, 50, and 100 *µ*m). We used the `annotation_25.nrrd` volume, which stores an integer brain-region identifier for each voxel coordinate (*x, y, z*) in the CCF. For a given CCF reference section, a two-dimensional annotation map was obtained by extracting the plane corresponding to the matched index (e.g., *z* = 205) from the annotation volume. Different colors were assigned to different region identifiers to generate a two-dimensional annotation image for visualization and downstream processing. Because the annotation volume (25 *µ*m isotropic resolution) differs from the pixel resolution of the CCF Nissl-stained images, a coordinate mapping between the Nissl-stained image and the annotation map is required. The regions of interest (ROIs) were independently extracted from the Nissl-stained image and the corresponding annotation image using Grounding DINO^39^. The Nissl ROI was resized to match the annotation ROI and then padded back to the original annotation image size, yielding a coordinate transformation between the two reference spaces. Anatomical annotation retrieval was performed in two steps. First, the experimental section was aligned with the corresponding Nissl-stained image. Second, the same coordinate transformation was applied to map the aligned experimental coordinates to the annotation space. Anatomical annotations were then assigned to the experimental spots.

#### Multi-omics spatial dataset of the mouse brain (MISAR-seq)

We used a multi-omics spatial dataset of the MISAR-seq containing paired spatial transcriptomics and ATAC-seq spatial sections of two samples. To increase the cross-modality discrepancy for validation, we introduced controlled perturbations to the Sample 2 section, generating a simulated Sample 2 slice. The spatial transcriptomics section (Sample 1) was used as a reference, while the spatial ATAC-seq section (Sample 2) served as a source. Each section was associated with a corresponding H&E-stained image. We first aligned the two H&E images to estimate a warping transformation. The resulting transformation was then applied to the spatial coordinates of all spots in the source section to achieve cross-modal alignment.

#### Mouse brain spatial tri-omics dataset

We further evaluated SPOmiAlign on a spatial tri-omics mouse brain dataset integrating MAGIC-seq spatial transcriptomics, MALDI imaging mass spectrometry-based metabolomics, and PLATO spatial proteomics. We first performed manual rotation and scaling between sections to obtain a reference alignment and then constructed SSIs for downstream comparison. For spatial transcriptomic SSIs, spot intensities were defined using UMI counts. For spatial metabolomic and proteomic SSIs, spot intensities were derived from measured metabolite and protein intensities, respectively. In the spatial transcriptomic and proteomic SSI, each spot was rendered as a square of 15 pixels, while in the spatial metabolomic SSI, a smaller square size of 12 pixels was used. All SSIs were constructed and visualized in Python using Matplotlib.

### Section matching

#### Robust dense feature matching

SPOmiAlign employs Robust Dense Feature Matching (RoMa)^24^ to compute dense pixel-level correspondences between two input sections. Both histological images and SSIs are supported as inputs. Given two images *A* (reference section) and *B* (source section), RoMa first encodes both images using a frozen DINOv2^40^ backbone, producing low-resolution yet highly robust coarse feature grids. For each coarse spatial location in the image *A*, a Gaussian Process (GP) matching module infers a global correspondence distribution over the coarse feature grid of the image *B*, capturing potential matching regions together with associated uncertainty. This probabilistic distribution is compactly encoded into a matching context representation. A Transformer-based decoder then maps this context to a discrete 64 × 64 anchor probability distribution *π*, indicating the most likely spatial region in image *B* that contains the correspondence, along with a matchability score *p*_*A*_ that estimates whether the location in image *A* is reliable for matching. The anchor distribution *π* is locally aggregated around its dominant mode using a weighted averaging scheme to generate an initial coarse warp defined in the coarse grid. Conditioned on this coarse warp, RoMa performs multi-scale coarse-to-fine refinement using high-resolution fine features, progressively correcting displacement estimates and updating matchability scores. The final output is a dense pixel-wise deformation field from image *A* to image *B*, together with per-pixel matchability maps *p*_*A*_ and *p*_*B*_, which can be directly used for image warping or for selecting reliable correspondence subsets.

#### Edge enhancement matching

For an input image of resolution 1152 × 864, RoMa generates dense correspondences for all 995,328 pixels. The original RoMa-based alignment strategy estimates the transformation using only the top *K* correspondences ranked by confidence (typically *K* = 5000). However, this strategy biases the selected correspondences toward highly textured interior regions, while tissue boundaries-often characterized by weaker visual features-are underrepresented, resulting in suboptimal alignment near tissue edges. To enhance boundary alignment while preserving global alignment performance, we propose an edge enhancement matching strategy that combines edge-aware confidence reweighting with spatial uniformization. To reduce the dominance of strong local textures and emphasize global structural features, we first apply Gaussian smoothing to the grayscale image:

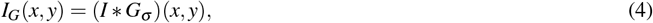

where *I*(*x, y*) denotes the original image and the Gaussian kernel *G*_*σ*_ is defined as

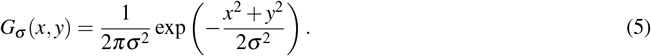

Following smoothing, Fourier-based edge enhancement is applied to detect prominent tissue boundaries. The Fourier transform of the smoothed image is given by

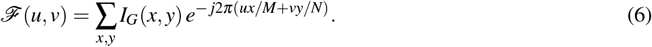

A high-pass filter *H*(*u, v*) is applied in the frequency domain to emphasize boundary-related components:

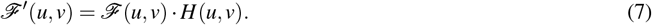

The enhanced image is obtained via an inverse Fourier transform:

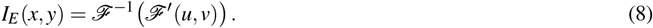

An edge mask is then generated by thresholding:

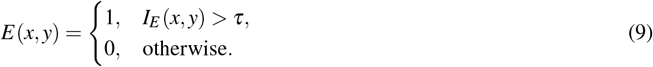

For each pair of matched pixels 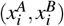, an edge-sensitive weight is computed as

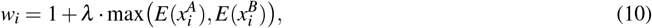

where *λ* controls the strength of the edge emphasis. Original RoMa matchability scores are reweighted as

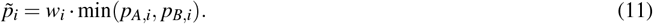

Although edge-aware reweighting improves boundary alignment, it can lead to excessive point density along tissue edges. To ensure a globally uniform correspondence distribution, we introduce a spatial uniformization module. The image domain is partitioned into sliding 3 × 3 pixel neighborhoods. Within each neighborhood Ω _*j*_, only the correspondence with the highest reweighted confidence is retained:

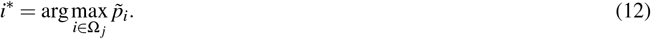

All correspondences remaining within the same neighborhood are discarded. The resulting correspondence subset is then used to estimate the transformation from image *A* to image *B*.

### Section warping

Depending on the spatial alignment scenario, different transformation models can be employed, including rigid, affine, and non-rigid transformations. SPOmiAlign adopts B-spline deformation for non-rigid alignment due to its flexibility and numerical stability. In this study, a two-stage alignment strategy is employed. An affine transformation is first estimated from the optimized correspondence subset to achieve coarse global alignment. This is followed by a non-rigid B-spline transformation for fine-scale refinement. The deformation field of the B-spline *T* (*x*) is defined as

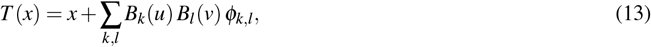

where *B*_*k*_(·) and *B*_*l*_(·) denote cubic B-spline basis functions and *φ*_*k,l*_ represent control point displacements. The estimated transformation is then applied to the spatial coordinates of all spots in the source section, producing warped coordinates that serve as the final aligned spatial coordinates for downstream analysis.

### Reassignment

Due to differences in sequencing technologies, spatial omics datasets often contain distinct numbers of spots and are measured at mismatched spatial resolutions, which preclude direct spot-wise correspondence and limit downstream multi-omic integration. To address this issue, we introduce a reassignment module to establish one-to-one correspondence between spots following section warping. After warping, two sections are designated as the reference section and the source section, where the section with higher spatial resolution is selected as the reference. Then, a nearest-neighbor interpolation strategy is applied. For each reference spot *i* with spatial coordinate (*x*_*i*_, *y*_*i*_), we identify its nearest neighboring spot *j* in the source section by minimizing the Euclidean distance

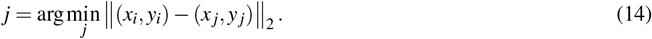

The molecular profile of the identified source point *j* is then assigned to the reference point *i*, generating reassigned profiles in the reference coordinate space. To correct for bias introduced by resolution differences—where a single source spot may be reassigned to multiple reference spots—we perform profile normalization. Let *n*_*j*_ denote the total number of times the profile of source spot *j* is assigned during reassignment. The final reassigned profile 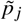 is calculated as

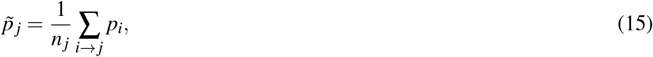

where the summation is taken over all reference spots *i* reassigned to the source spot *j*, and *p*_*i*_ denotes the molecular profile associated with the spot *i*. In practice, spatial regions captured by different omics may not fully overlap. To avoid erroneous reassignment in non-overlapping regions, a distance-based filtering criterion is introduced. We first compute the average nearest-neighbor distance among all spots within the reference and source sections, denoted *D*_*r*_ and *D*_*s*_, respectively. Then, a global distance scale *D* is defined as

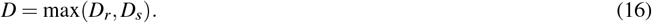

Based on this scale, a distance threshold *K* is defined as

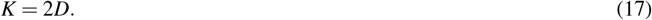

For each reference spot *i*, a reassignment is performed only if the distance to its nearest neighboring source spot satisfies

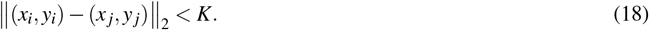

Reference spots without a valid neighboring source spot (distance ≥ *K*) are excluded from reassignment and omitted from subsequent integrative analyses.

### Vertical multimodal integration

After section matching and reassignment, multimodal sections were brought into one-to-one spot correspondence, yielding a unified set of spatial locations shared across modalities. Vertical multimodal integration was performed using SpatialGLUE^29^ to jointly embed multimodal molecular profiles into a shared latent space. Subsequently, Leiden clustering was applied to the integrated embeddings to identify spatial domains. To determine the optimal clustering resolution, we evaluated clustering hierarchies using Clustree^30^, which enables visualization of cluster stability across resolutions. Based on Clustree analysis, the optimal number of clusters was selected for spatial domain identification and downstream refinement.

### Differential Analysis of Multi-omics Data

Differential analyses of spatial transcriptomics (ST), spatial proteomics (SP), and spatial metabolomics (SM) were performed between the inner molecular layer (IML) and outer molecular layer (OML) subregions identified within 5 clusters. For ST and SP, normalized gene expression and protein abundance matrices were extracted from the integrated AnnData object. Differential testing was conducted using the Wilcoxon rank-sum test (implemented in Scanpy) to ensure methodological consistency ^41^. Log-fold changes (logFC) were calculated based on the difference in mean expression or abundance between subregions. For SM, normalized metabolite intensity profiles were compared using the Wilcoxon rank-sum test to account for non-Gaussian signal distributions. For all modalities, P-values were adjusted for multiple testing using the Benjamini–Hochberg (BH) procedure. Features with an adjusted P-value < 0.1 and logFC > 0.25 were considered statistically significant. Differentially upregulated and downregulated features were extracted for downstream integration.

### Metabolite Annotation

Differentially abundant spatial metabolomic features were annotated by matching observed m/z values against the Human Metabolome Database (HMDB)^42,43^. Matching was performed within a mass tolerance window of 3 ppm. Putative identities were assigned based on molecular formula and monoisotopic mass. To improve coverage, multiple adduct forms in negative ionization mode were considered, including [M-H]-, [M+Cl]-, [M+FA-H]-, and [M+Ac-H]-. Annotated metabolites were classified into superclass, class, and subclass categories according to HMDB taxonomy and grouped into lipid and non-lipid categories for pathway interpretation.

### Integrated analysis of ST, SP, and SM

#### ST-SP integration

We adopted a stepwise integration strategy centered on transcription–protein concordance to identify molecular programs supported across modalities. To identify robust molecular programs supported across transcriptional, proteomic, and metabolic layers, we adopted a stepwise integration strategy centered on the concordance between Spatially Variable Genes (SVGs) and Spatially Variable Proteins (SVPs). ST and SP datasets were first aligned based on shared gene symbols, and independent differential analyses were performed to identify features regulated between the inner (IML) and outer (OML) molecular layers. We initially prioritized gene–protein pairs exhibiting significant positive associations using Spearman’s rank correlation analysis (Spearman’s ρ>0.1, FDR <0.05). To further ensure robust regulatory consistency, a coherence filtering threshold of 0.45 was applied to retain features exhibiting strong directional agreement, thereby defining a high-confidence “ST–SP concordant signature.” These features were ranked according to their combined differential signals and visualized via heatmaps.

#### ST–SP–SM Integration

Building upon this validated transcriptomic–proteomic axis, spatial metabolomics (SM) data were subsequently integrated to characterize the specific metabolic states and lipid microenvironments associated with these coordinated molecular programs. Spatial spots were ordered based on the ranked expression scores of the ST–SP concordant signature. Normalized profiles for genes, proteins, and metabolites were extracted from spatially aligned coordinates. Spearman’s rank correlation analysis was performed to identify metabolite–gene and metabolite–protein pairs exhibiting concordant spatial variation (Spearman’s ρ > 0.2, FDR < 0.05). Significant associations were summarized to define metabolite classes preferentially enriched in transcriptionally and proteomically defined spatial domains (IML vs. OML).

### Functional enrichment analysis

Functional enrichment was performed to interpret biological programs driving sublayer stratification. Differentially expressed genes (ST) and proteins (SP) were analyzed independently using Enrichr^44^. Queries included Gene Ontology (GO) Biological Process (2025), Reactome Pathways (2024), SynGO (2024), and MGI Mammalian Phenotype (Level 4, 2024), alongside brain cell-type reference datasets including the Allen Brain Atlas (10x scRNA 2021) and Azimuth (2023). Enrichment analysis was further applied to the ST–SP concordant gene set to identify pathways robustly supported by both transcriptional and proteomic layers. Statistical significance was assessed using Fisher’s exact test with Benjamini–Hochberg correction. Terms with an adjusted P-value < 0.05 were considered significant. Results were visualized using ranked bar plots to highlight consistent biological processes.

### Metrics for evaluating registration to the CCF

#### Full width at half maximum (FWHM)

To assess spatial registration precision at the molecular level, we quantified the full width at half maximum (FWHM) of marker-specific spatial distributions after alignment to the Allen Common Coordinate Framework (CCF). For a given molecular marker, we first estimate its spatial density across all spots in the aligned section. For each spot, we then computed the shortest Euclidean distance to the boundary of the corresponding CCF anatomical region obtained after annotation transfer. The distribution of these distances was summarized as a one-dimensional density function. Let *d* denote the distance to the boundary of the corresponding CCF region, and let *f* (*d*) denote the estimated density function of *d*. The FWHM was defined as the width of the distance distribution at half of its maximum density value, that is,

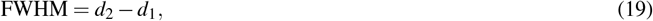

where *d*_1_ and *d*_2_ satisfy

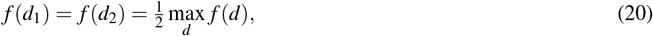

with *d*_1_ *< d*_2_. A smaller FWHM indicates that marker-positive spots are more closely localized around the corresponding anatomical boundaries of the CCF. reflecting the higher spatial registration accuracy between the spatial omics data and the CCF.

#### Dice similarity coefficient

To evaluate region-level alignment accuracy, we computed the Dice similarity coefficient between manually annotated brain regions in the spatial omics data and the corresponding brain regions automatically assigned by SPOmiAlign after CCF-based annotation retrieval. For a given brain region, let *A* denote the set of spots labeled by manual annotation and *B* denote the set of spots assigned to the corresponding CCF region by SPOmiAlign. The Dice similarity coefficient is defined as

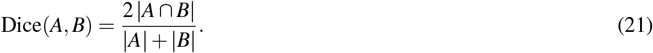

A larger Dice coefficient indicates greater overlap between ground-truth annotations and CCF-retrieved annotations, corresponding to higher registration accuracy.

### Metrics for evaluating multimodal alignment

#### Cell type-based Pearson correlation coefficient and cosine similarity

In the evaluated datasets, Cell type annotations were available for section S1 but absent for section S2. To enable a cell type–based alignment assessment, section S2 was manually annotated. To ensure consistency of cell-type definitions across sections, annotations in S1 were further curated by merging fine-grained subclasses into broader categories shared between S1 and S2. For each matched cell type, we assessed the spatial compositional correspondence between modalities after alignment. Specifically, the Pearson correlation coefficient (PCC) and cosine similarity were calculated between the representations of the cell-type in the matched spots in S1 and the aligned spots in S2. Higher PCC and cosine similarity values indicate stronger agreement in the spatial distributions of the cell-type between modalities, reflecting improved multimodal alignment.

#### Bivariate Moran’s *I* for spatial co-localization

In spatial multi-omics data, different modalities are characterized by distinct molecular measurements, such as gene expression levels in spatial transcriptomics and metabolite intensities in spatial metabolomics. To evaluate multimodal alignment, it is therefore essential to quantify the spatial concordance of co-localized biomarkers across modalities. Previous single-modality studies commonly employ univariate Moran’s *I* to measure spatial autocorrelation of molecular features. In the multimodal setting, we extend this concept by computing bivariate Moran’s *I*, which captures the spatial association between biomarkers from two different modalities. Formally, let *x*_*i*_ and *y* _*j*_ denote the values of two biomarkers from different modalities in spatial locations *i* and *j*, respectively, and let *w*_*ij*_ denote the spatial weight matrix encoding neighborhood relationships. The bivariate Moran’s *I* is defined as

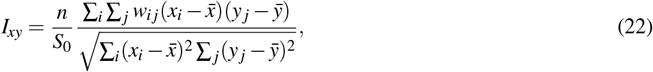

where *n* is the total number of spatial locations, 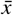 and 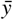 denote the mean values of the two biomarkers, and

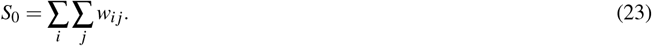

A higher value of the bivariate Moran’s *I* indicates stronger spatial co-localization between biomarkers across modalities, corresponding to a more accurate multimodal alignment.

### Metrics for evaluating multi-omic integration and clustering

In addition, clustering performance was evaluated using standard clustering metrics, including adjusted Rand index (ARI), normalized mutual information (NMI), silhouette score, Calinski–Harabasz index, homogeneity, completeness, and V-measure. These metrics were calculated based on spot-matched labels after alignment and reassignment.

## Data availability

All datasets used in this study are publicly available. The Slide-seq dataset, a high-throughput sequencing-based spatial transcriptomics technology with near-cellular resolution comprising 101 adult mouse brain coronal sections that span the entire anteroposterior axis, was obtained from an online resource at https://docs.braincelldata.org/downloads/index.html/Slide-seq_Data. The MERFISH dataset, a high-throughput imaging-based spatial transcriptomics technology with single-cell resolution consisting of 25 adult mouse brain sagittal sections, was obtained from the Allen Brain Cell Atlas at https://alleninstitute.github.io/abc_atlas_access/descriptions/Zhuang-ABCA-3.html. The Allen Common Coordinate Framework (CCF) and the corresponding brain region annotations were obtained from the Allen Brain Atlas. Nissl-stained reference images of the adult mouse brain CCF used for alignment are publicly available for coronal views at https://mouse.brain-map.org/experiment/thumbnails/100048576?image_type=atlas and sagittal views at https://mouse.brain-map.org/experiment/thumbnails/100042147?image_type=atlas. Brain region annotations corresponding to the 25 *µ*m resolution CCF were downloaded from the file `annotation_25.nrrd` at https://download.alleninstitute.org/informatics-archive/current-release/mouse_ccf/annotation/ccf_2017/annotation_25.nrrd. The correspondence between the brain region identifiers and the anatomical labels was obtained from `structure _tree _safe _2017.csv`, available at https://github.com/cortex-lab/allenCCF/blob/master/structure_tree_safe_2017.csv. The MISAR-seq dataset, a spatial multi-omics profiling dataset of the mouse brain that jointly measures chromatin accessibility and gene expression, was obtained from a public repository in https://github.com/gpenglab/MISAR-seq/blob/main/Data/Download. The spatial multi-omics mouse brain dataset that integrates MALDI imaging mass spectrometry-based metabolomics, MAGIC-seq spatial transcriptomics, and PLATO spatial proteomics was obtained from a public repository at https://github.com/bioinfo-biols/Flow2Spatial/tree/main/datasets. Source data are provided with this paper.

## Code availability

The code for the SPOmiAlign program is available on GitHub at https://github.com/wangyiyuyang/SPOmiAlign.

## Acknowledgements

The authors thank Prof. Ning Shen for helpful advice on this study. The authors thank Yicheng Liu, Zhiyu Wen, and Ningyuan You for their assistance in data collection. This work was supported by the Yuhang District Postdoctoral Research Funding Program, Hangzhou, Zhejiang Province, China. A subset of images was created with BioRender.com, with permission.

## Author contributions statement

Y.W. conceived, designed, and supervised the study, designed the model, collected data, performed data analysis, and drafted the original manuscript; Z.H. designed the multi-omics study, collected data, performed data analysis, drafted the original manuscript, and revised the manuscript. Y.Y. performed the data analysis and developed the code. All authors read and approved the manuscript.

## Competing interests

The authors declare no competing interests.

## Notes

https://github.com/wangyiyuyang/SPOmiAlign

### Competing Interest Statement

The authors have declared no competing interest.

