## supplemental figures for "SPOmiAlign: A modality-agnostic framework for robust and scalable spatial multimodal alignment via feature matching"

### Supplementary figures

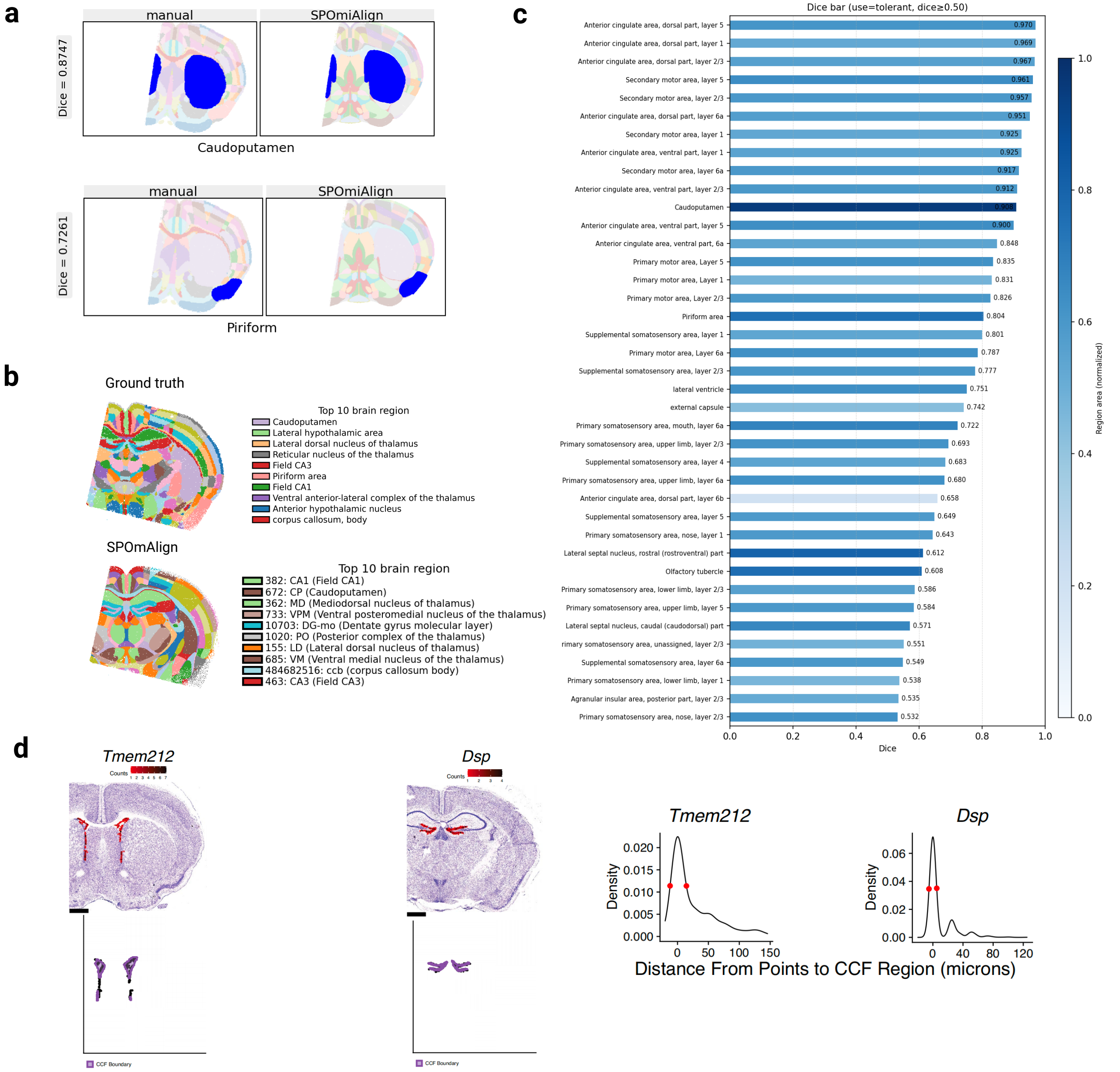


**Supplementary Figure 1: Additional evaluation of anatomical annotation accuracy on Slide-seq mouse brain sections. a** Dice similarity coefficients for two representative corresponding brain regions between SPOmiAlign-derived annotations and ground truth in Slide-seq mouse brain coronal section ID 29. **b** Comparison of anatomical annotations assigned by SPOmiAlign and ground truth across all brain regions in Slide-seq coronal section ID 43, with the top ten regions ranked by area shown in the legend. **c** Dice similarity coefficients for all corresponding brain regions with Dice $> 0.5$ between SPOmiAlign and ground truth annotations in Slide-seq coronal section ID 29; color indicates region area. **d** Distance density distribution between ground-truth enriched gene expression spots and corresponding CCF anatomical boundaries.

| Dataset | Mouse brain spatial two−omic | Mouse brain spatial three−omic | Mouse brain 3D sagittal | Mouse brain 3D cornal | Mouse brain 3D cornal |
|---|---|---|---|---|---|
| Modality | Spatial transcriptomic、Spatial ATAC−seq | Spatial transcriptomic、Spatial proteomic、Spatial metabolomic | Spatial transcriptomic | Spatial transcriptomic | Spatial transcriptomic |
| Technology | MISAR−seq | MAGIC−seq、PLATO、MALDI−MSI | MERFISH | Slide−seq | Slide−seq |
| Spot number | 1939 | 3908 | 92497 | 169683 | 213885 |
| runtime1(s) | 8.4193 | 9.7285 | 12.1604 | 11.3602 | 11.2004 |
| runtime2(s) | 6.1438 | 9.7763 | 11.6144 | 11.4038 | 11.7882 |
| runtime3(s) | 5.0793 | 9.5984 | 11.7221 | 11.6043 | 11.2525 |
| runtime4(s) | 5.2141 | 6.6163 | 11.5386 | 11.5264 | 11.3965 |
| runtime5(s) | 4.4901 | 8.7012 | 16.4682 | 11.2858 | 11.3575 |

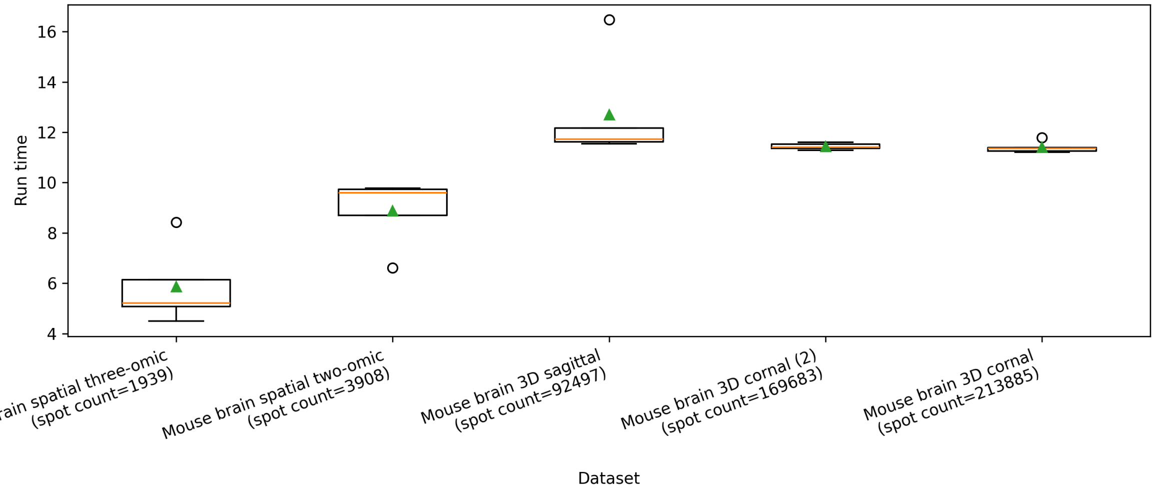


**Supplementary Figure 2: Box plots of SPOmiAlign runtime from five repeated runs across five datasets, ordered by increasing spot count.**

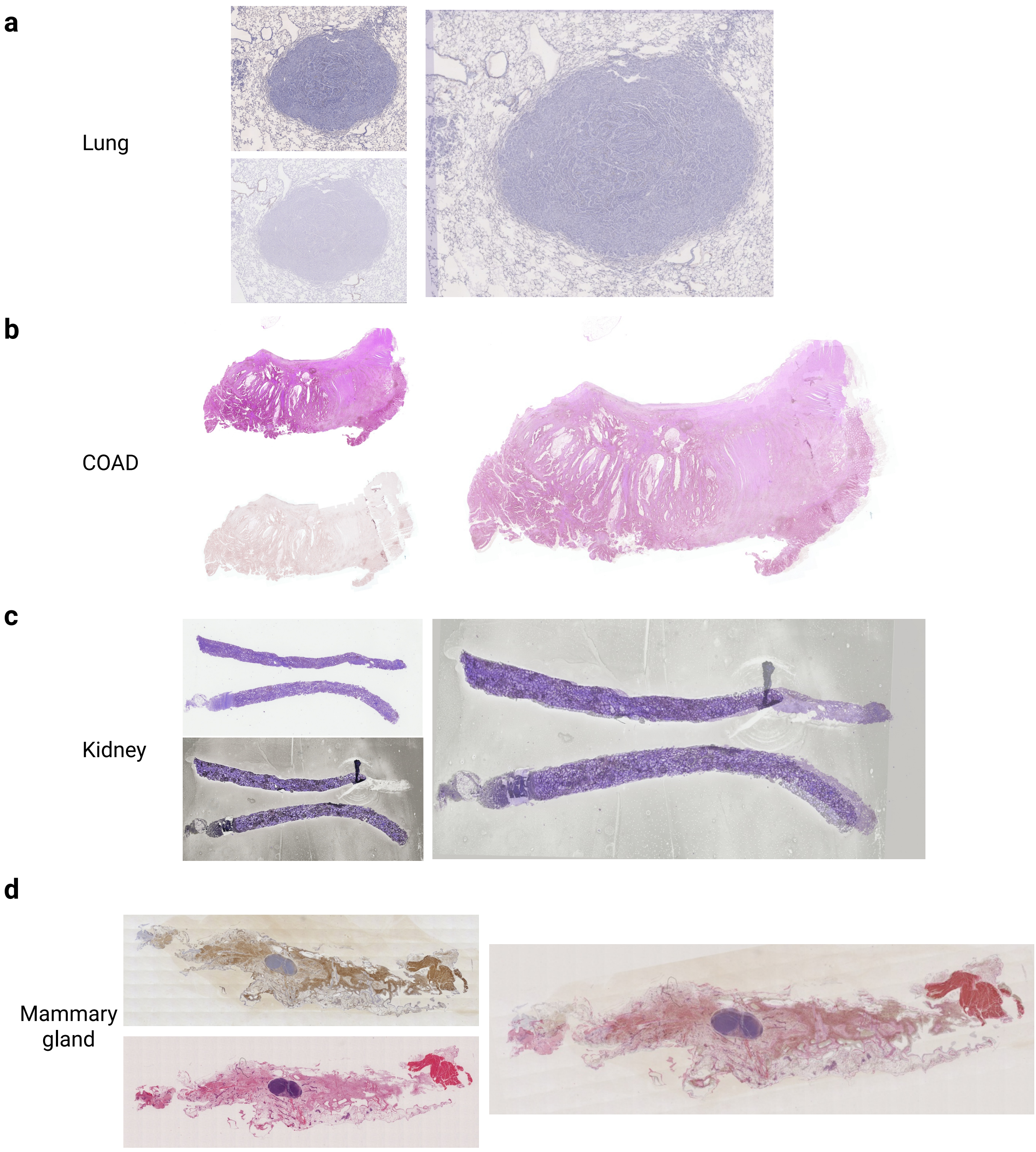


**Supplementary Figure 3: Registration of pathological whole-slide images across serial sections from the same tissue in four diseases. a–d** The two panels on the left show the original, unaligned whole-slide images, while the right panel shows the overlaid images after alignment.

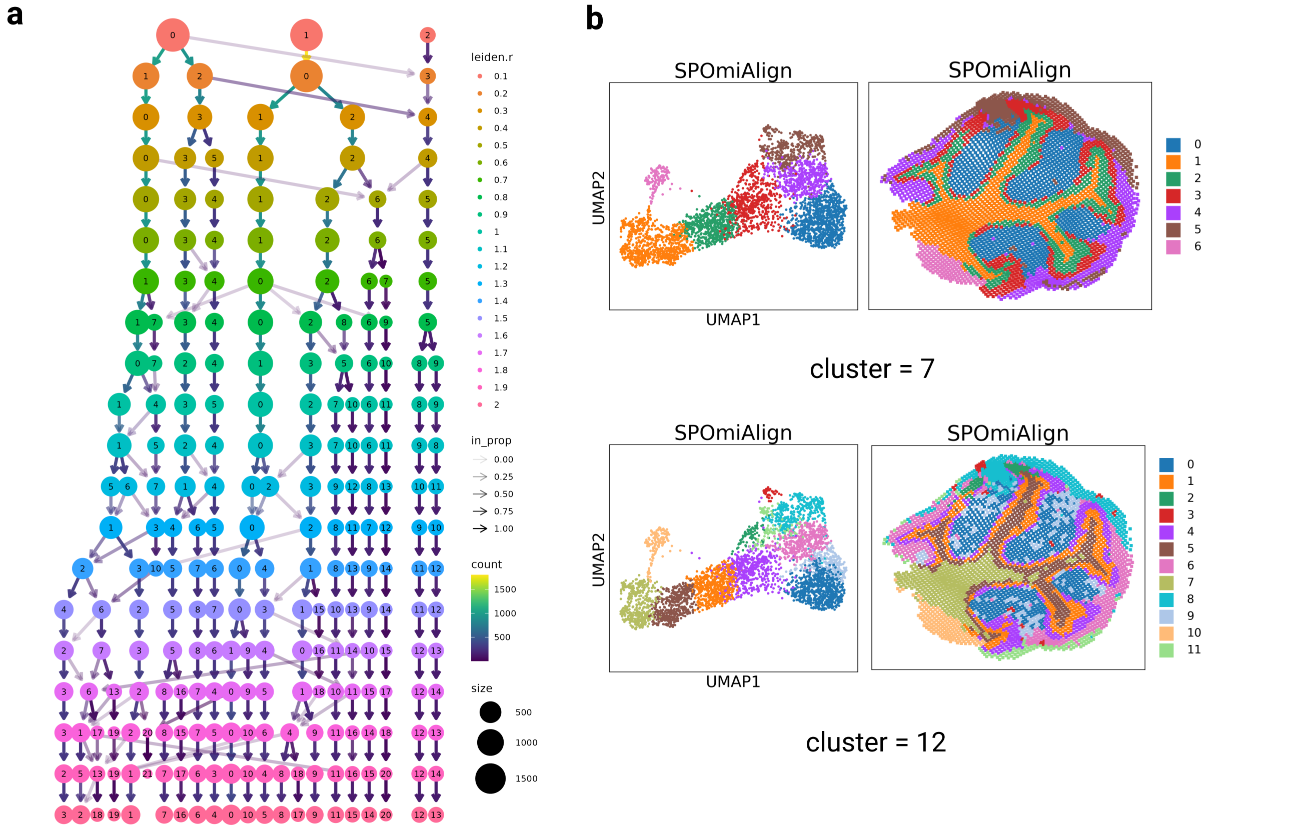


**Supplementary Figure 4: Clustree-based assessment of spatial domain stability in integrated tri-omics data. a** Clustree representation of clustering hierarchies constructed from the integrated spatial tri-omics dataset. **b** Spatial domain in two relatively stable clustering configurations (clusters = 7 and 12) revealed by clustree analysis.

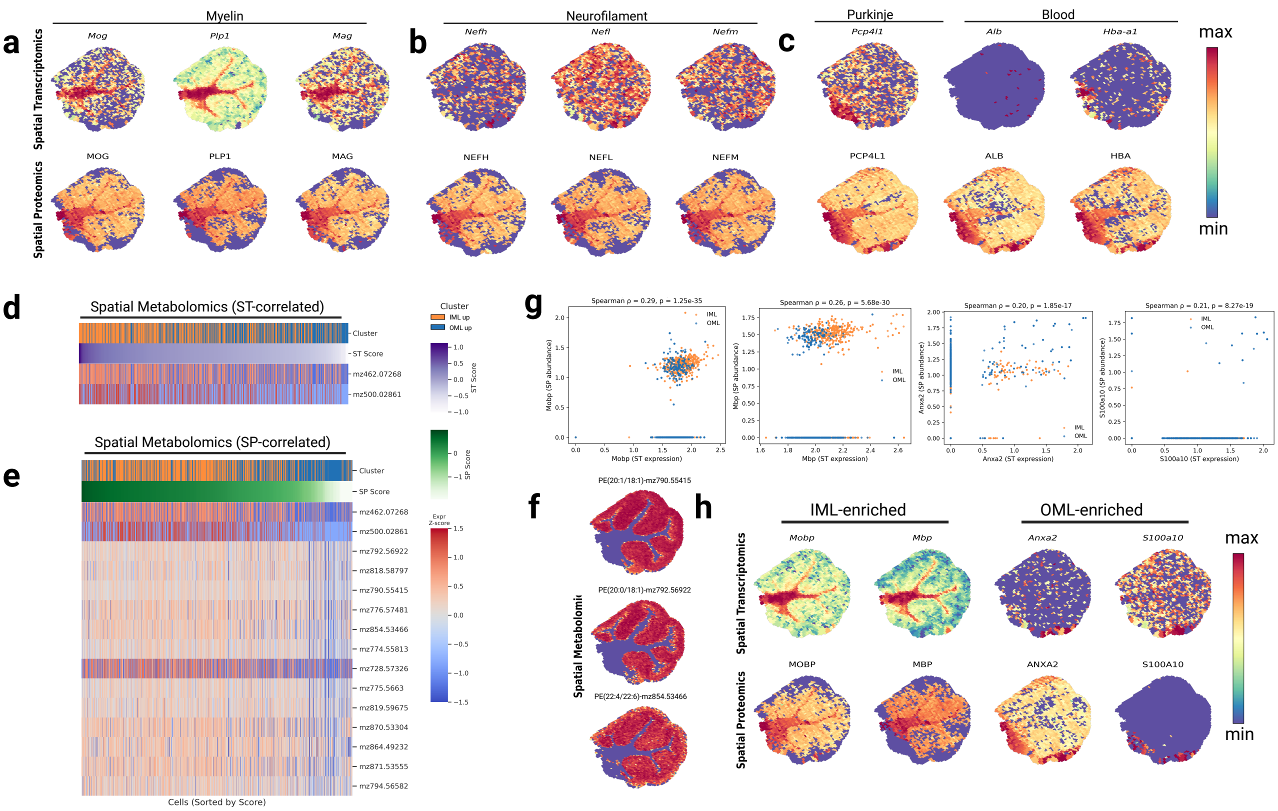


**Supplementary Figure 5: Integrated multi-omics characterization of molecular layer substructures. a–c** Spatial distribution maps of representative features identified from integrated multi-omics analyses. **a** Myelin- and oligodendrocyte-related genes (*Mog*, *Plp1*, *Mag*) enriched in the inner molecular layer (IML) relative to the outer molecular layer (OML). **b** Neurofilament-related genes (*Nefh*, *Nefl*, *Nefm*) enriched in the IML relative to the OML. **c** Purkinje cell- and blood-related proteins enriched in the OML relative to the IML. **d,e** Heatmaps of non-annotated metabolites identified via correlation analysis with spatial transcriptomics (ST) (**d**) and spatial proteomics (SP) (**e**) based on SVG–SVP pairs. The heatmaps display metabolites positively associated with gene expression and protein abundance patterns (Spearman correlation $> 0.2$, FDR $< 0.05$). **f** Spatial distribution maps of PE metabolites identified in Fig. 5h. **g** Spearman correlation analysis of SVG–SVP pairs for featured genes. **h** Spatial distribution maps of representative transcripts and proteins identified from integrated multi-omics analyses, demonstrating their concordant localization within molecular layer substructures. IML-enriched features (left), OML-enriched features (right).
